# Late ESCRT Machinery Mediates The Recycling And Rescue of Invariant Surface Glycoprotein 65 in *Trypanosoma brucei*

**DOI:** 10.1101/2020.06.12.146795

**Authors:** Khan Umaer, James D. Bangs

**Author notes:** Corresponding Author: James D. Bangs.

## Abstract

The <u>E</u>ndosomal <u>S</u>orting <u>C</u>omplex <u>R</u>equired for <u>T</u>ransport machinery consists of four protein complexes (ESCRT 0-IV) and the post ESCRT ATPase Vps4. ESCRT mediates cargo delivery for lysosomal degradation via formation of multivesicular bodies. *Trypanosoma brucei* contains orthologues of ESCRT I-III and Vps4. Trypanosomes also have a ubiquitinylated invariant surface glycoprotein (ISG65) that is delivered to the lysosome by ESCRT, however, we previously implicated TbVps4 in rescue and recycling of ISG65. Here we use conditional silencing to investigate the role of TbVps24, a phosphoinositide-binding ESCRT III component, on protein trafficking. TbVps24 localizes to the TbRab7^+^ late endosome, and binds PI(3,5)P_2_, the product of the TbFab1 kinase, both of which also localize to late endosomes. TbVps24 silencing is lethal, and negatively affects biosynthetic trafficking of the lysosomal markers p67 and TbCathepsin L. However, the major phenotype of silencing is accelerated degradation and depletion of the surface pool of ISG65. Thus, TbVps24 silencing phenocopies that of TbVps4 in regard to ISG65 trafficking. This presents a paradox since we have previously found that depletion of TbFab1 completely blocks ISG65 turnover. We propose a model in which late ESCRT components operate at two sites, one PI(3,5)P_2_-dependent (degradation) and one PI(3,5)P_2_-independent (recycling), to regulate ISG65 homeostasis.

## INTRODUCTION

*Trypanosoma brucei* (African trypanosomes) is a protozoan parasite that causes human African trypanosomiasis and related diseases in livestock; it is transmitted by an insect vector, tsetse flies. As an early diverging eukaryote, it contains most of the typical eukaryotic secretory and endocytic organelles, including the endoplasmic reticulum (ER), ER exit sites, the Golgi, a single terminal lysosome and the core eukaryotic endosomal compartments (early, late and recycling). However, the reduced copy number of these organelles, and the small size (~25 x 3 μm) and highly polarized morphology of trypanosomes, suggests streamlining for efficient secretory and endocytic transport (Bangs, 2011; Engstler, Bangs, & Field, 2006; Overath & Engstler, 2004; Silverman & Bangs, 2012). All endocytosis and secretion in bloodstream form (BSF) trypanosomes proceeds via a specialized invagination of the plasma membrane termed the flagellar pocket. All the other organelles that participate in endocytosis and secretion are located between the nucleus (positioned centrally) and the flagellar pocket (positioned towards the posterior end of the cell). Two vesicular pathways deliver cargo of different origins to the single terminal lysosome: the endocytic pathway delivers nutritional cargo taken up via the flagellar pocket, and the biosynthetic pathway delivers newly synthesized proteins from the Golgi apparatus. In addition, internalized surface components, such as variant surface glycoprotein (VSG), transferrin receptor (TfR), and invariant surface glycoproteins (ISGs), can be sorted from lysosomal cargo in endosomes and recycled to the cell surface via the flagellar pocket (Engstler et al., 2004; Kabiri & Sterverding, 2000; Leung, Riley, Carrington, & Field, 2011). There are several points at which these routes intersect, in particular the late endosome/multivesicular body (LE/MVB), where biosynthetic and endocytic cargo meet en route to the lysosome. As in other eukaryotes the LE/MVB in trypanosomes is marked by the small GTPase TbRab7, as well as by conserved components of the ESCRT (<u>e</u>ndo<u>s</u>omal <u>c</u>omplex <u>r</u>equired for <u>t</u>ransport) machinery (Leung, Dacks, & Field, 2008; Silverman, Muratore, & Bangs, 2013; Silverman, Schwartz, Hajduk, & Bangs, 2011).

Eukaryotic MVBs are large specialized LEs containing intralumenal vesicles (ILVs) that are formed by internal reverse-topological budding of the endosome membrane (Hurley, 2015; Schöneberg, Lee, Iwasa, & Hurley, 2017; Vietra, Radulovic, & Stenmark, 2020). Membrane deformation and budding are mediated by the sequential actions of the cytoplasmic ESCRT 0, I and II multiprotein complexes; membrane scission is mediated by the ESCRT III complex and Vps4 ATPase; and the residual ESCRT proteins are then disassembled and recycled by Vps4. The ESCRT machinery, particularly ESCRT III and Vps4, functions in many other cellular processes besides MVB formation, including exosome biogenesis, cellular abscission, viral budding, plasma membrane repair, nuclear envelope maintenance, and more. However, the function most relevant for this work is the recruitment of ubiquitinylated transmembrane proteins by ESCRT 0 and I proteins for inclusion in ILVs and subsequent degradation following MVB:lysosomal fusion.

Trypanosomes do not have morphologically obvious MVBs, although atypical MVB-like structures have been detected in cells stressed by knockdown of proteins of the endocytic and secretory pathway (Allen, Liao, Chung, & Field, 2007; Peck et al., 2008). Nevertheless, with the exception of ESCRT 0, they have orthologues of all the major components of the other ESCRT complexes (Leung et al., 2008). Of these, only TbVps23 (ESCRT I) and TbVps4 have been characterized in detail (Leung et al., 2008; Silverman et al., 2013). Both co-localize with the LE marker TbRab7 suggesting MVB functionalities (Silverman et al., 2013). Knockdown of TbVps4 had varying negative effects on both biosynthetic and endocytic lysosomal trafficking, but its most prominent effect was on turnover of ISG65 (invariant surface glycoprotein 65 kDa), a type I transmembrane glycoprotein of unknown function. Internalization, recycling, and ultimate lysosomal degradation of ISG65 is regulated by ubiquitinylation of lysine residues in the C-terminal cytoplasmic domain (Chung, Carrington, & Field, 2004; Chung, Leung, Carrington, & Field, 2008; Leung et al., 2011). TbVps4 knockdown, which shuts down the entire ESCRT pathway, dramatically increased lysosomal degradation of ISG65, and this phenotype is recapitulated by knockdown of TbRab7 (Silverman et al., 2013). These data strongly suggest that the trypanosomal ESCRT machinery is involved in rescuing ISG65 from lysosomal targeting by recycling to the cell surface from the late endosome (but see Discussion). Although the ESCRT machinery is typically involved in recruitment of ubiquitinylated cargo for degradation, a role in recycling of cell surface proteins is not unprecedented—ESCRT machinery, specifically the ESCRT III component Vps24, is involved in recycling of both epidermal growth factor receptor (EGFR) and claudin I in mammalian cells (Baldys & Raymond, 2009; Dukes et al., 2011).

Phosphoinositides (PIs) are low abundance membrane phospholipids that mediate critical cellular functions in membrane trafficking (Balla, 2013; Di Paolo & De Camilli, 2006). Operating as spatiotemporally controlled signposts for compartmental membrane identity (Behnia & Munro, 2005), they mediate the reversible recruitment of cytosolic proteins and protein complexes to specific membranes at specific time points. Specific PI kinases and phosphatases can interconvert the phosphorylation status of the 3-, 4-, and 5-OH groups in the inositol ring of phosphatidylinositol singularly or in combination to produce seven distinct PIs (Balla, 2013). Two of these play roles in recruitment of the ESCRT machinery to nascent MVBs, formation of which likely starts at the early or sorting endosome. PI3P, the product of the PI-3 kinase Vps34, indirectly recruits ESCRT 0 and ESCRT II to budding sites (Balla, 2013). PI(3,5)P_2_, the product of the PI(3)P-5 kinase Fab1 localizes to the late endosome, where it binds directly to the mammalian ESCRT III component Vps24 (aka CHMP3), and to its paralogues CHMP2a and CHMP4b (Catimel et al., 2008; Whitley et al., 2003). Thus, PI(3,5)P_2_ is directly implicated in recruitment of ESCRT III to the nascent MVB. It should be noted that Vps24 also binds *in vitro* to PI(3,4)P_2_, but not to other phosphoinositides (Whitley et al., 2003).

We have previously investigated the function of the trypanosomal orthologue, TbFab1, and its product PI(3,5)P_2_ in endo/lysosomal trafficking (Gilden et al., 2017). Both localize prominently to TbRab7-positive LE and the TbCatL (lumenal protease)-positive terminal lysosome. Knockdown of TbFab1 reduced PI(3,5)P_2_ levels, but had minimal effects on biosynthetic and endocytic trafficking to the lysosome. Startlingly however, TbFab1 ablation resulted in complete blockage of ISG65 turnover in the lysosome. To investigate this phenomenon we now turn our attention to TbVps24 because of its phosphoinositide binding properties. In this study, utilizing conditional RNAi knockdown methodology, we take a more detailed look at the functions of TbVps24 in bloodstream form *T. brucei* with an emphasis on its association with phosphoinositides, and its role in ISG65 homeostasis. Our results represent a first detailed analysis of Vps24 functions in trypanosomes.

## Results

### TbVps24 is essential for bloodstream form *T. brucei* survival

We created a BSF cell line with an inducible stem loop dsRNA vector targeting the endogenous *TbVps24* gene. Following induction with tetracycline, the steady state levels of TbVps24 mRNA was reduced by 69.62 ± 3% (n=3) at 12 hr and 61.18 ± 4% (n=3) at 16 hr, whereas cell growth slowed after 12 hr with complete cessation at 16 hr and cell death following rapidly thereafter (Figs. 1A & 1B). However, the steady state localizations of BiP and p67 showed no disruption of internal ER or lysosomal morphology, respectively at 16 hr post-induction as determined by fluorescence microscopy (Fig. 1C). All subsequent experiments were performed at 12 hours of induction, at the point where TbVps24 knockdown first impacts growth without affecting gross and internal morphology.

**Figure 1.**
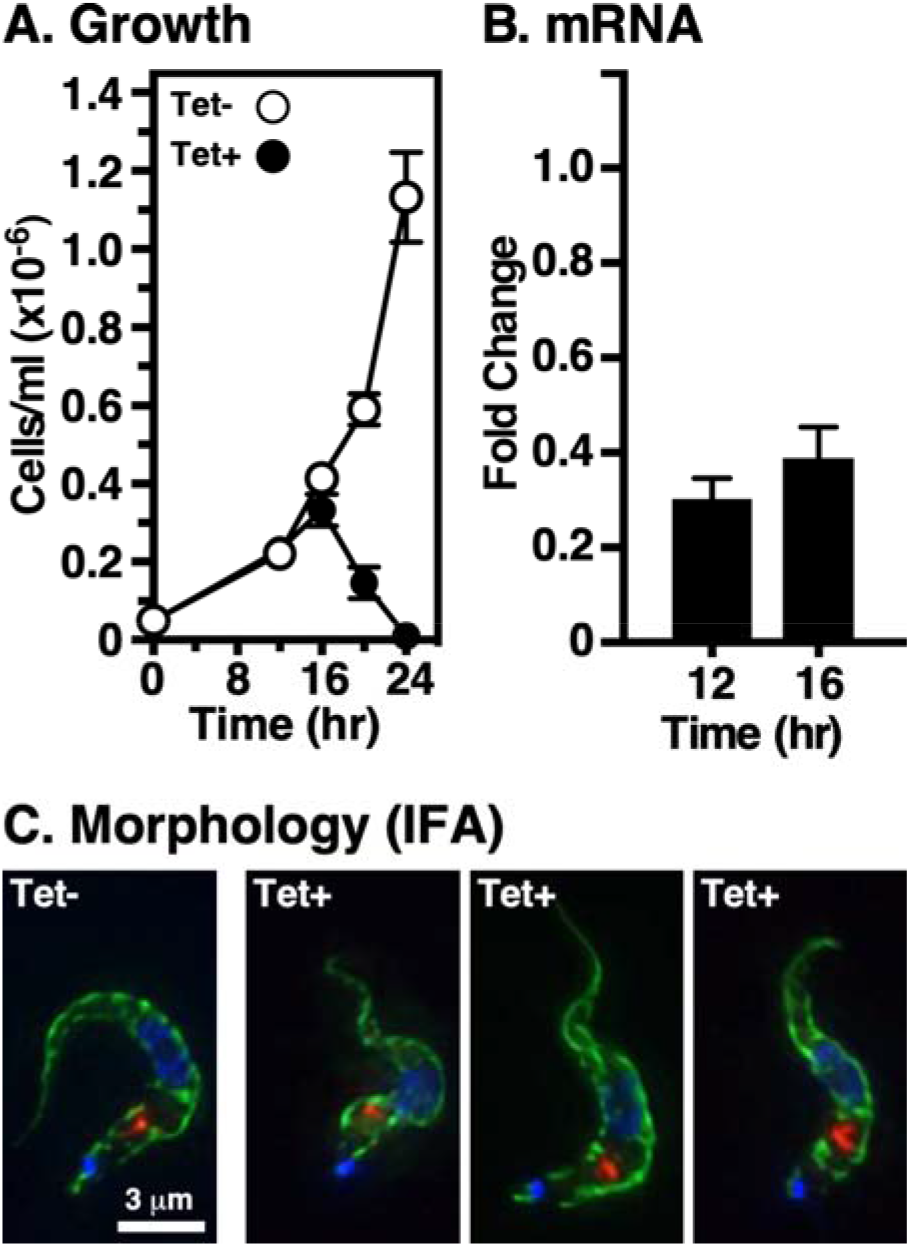
Vps24 is essential for proliferation of bloodstream *T. brucei* cells. The TbVps24 RNAi cell line was grown in the presence of tetracycline (1 μg/ml) to induce specific knockdown. **A**. Cell density was measured and plotted against time (mean ± std. dev., n=3 biological replicates). **B.** The extent of TbVps24 mRNA knockdown was assayed by real-time qRT-PCR at the indicated time points. Values are normalized to uninduced controls (mean ± std. dev., n=3 biological replicates). All subsequent RNAi experiments (except IFA, next panel) were performed at 12 hr of silencing. **C.** Immunofluorescence microscopy was performed with fixed permeabilized control and silenced (16 hr) cells using anti-BiP (green) and anti-p67 (red) to localize the ER and lysosomes respectively. Nuclei and kinetoplasts were stained with DAPI (blue). Representative 3-channel summed stack projections are presented. Bar indicates 3 μm.

### Localization of TbVps24

Previous work in our lab, and others, has localized TbVps23, TbVps28 and TbVps4 predominantly to the LE in close proximity to the lysosome (Leung et al., 2008; Silverman et al., 2013). All three proteins also showed diffuse cytoplasmic staining consistent with a cytoplasmic pool, as expected for the ESCRT machinery. In order to localize TbVps24, we created a BSF cell line expressing C-terminal HA-tagged TbVps24 and N-terminal Ty-tagged TbRab7. TbVps24-HA staining was mainly observed in the post-nuclear region with prominent co-localization with the LE marker TbRab7 (Figure 2A), and closely juxtaposed to the lysosomal membrane marker p67 (Figure 2B). However, we also observed diffuse cytoplasmic TbVps24 staining not associated specifically with the late endosome or lysosome, consistent with both free cytoplasmic and membrane-bound pools. These results are consistent with prior localizations of TbVps23, TbVps28 and TbVps4 (Leung et al., 2008; Silverman et al., 2013).

**Figure 2.**
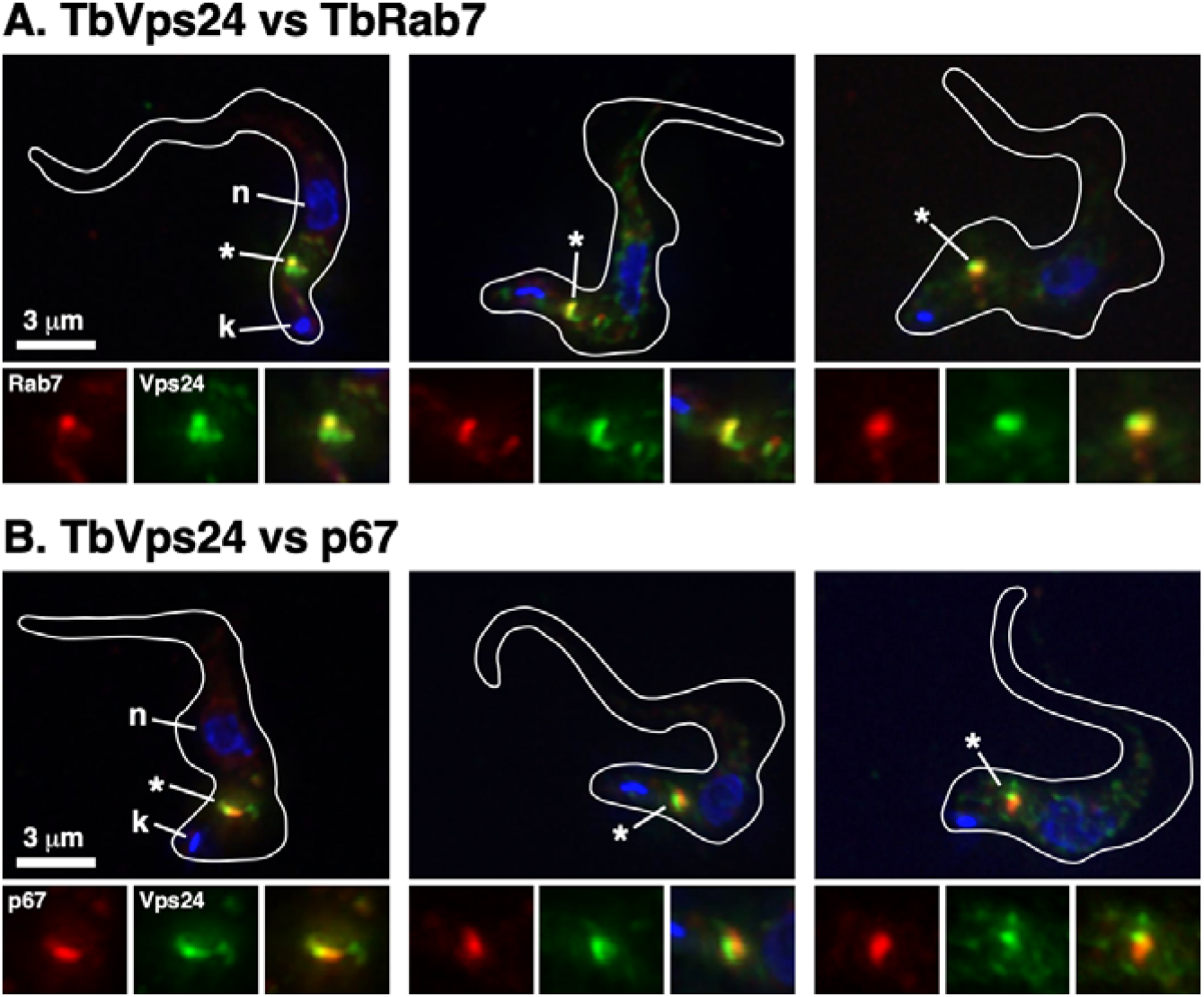
Localization of TbVps24. A double tagged TbVps24:HA/Ty:TbRab7 cell line was used for immunolocalization. Permeabilized cells were stained with anti-HA (green) and anti-Ty (red) to localize relative to the Rab7+ LE (**A**), or anti-HA (green) and anti-p67 (red) to localize relative to the terminal lysosome (**B**). Three-channel summed stack projections are presented. Cell outlines traced from matched DIC images; kinetoplasts (k) and nuclei (n) are indicated. The LE/lysosomal regions (asterisk) are presented in 1.5x magnified single- and three-channel format at the bottom of each image.

### TbVps24 silencing affects biosynthetic lysosomal trafficking

To investigate the role of TbVps24 in trafficking of newly synthesized proteins to the lysosome, we used TbCatL, a soluble lysosomal thiol protease, and p67, a lysosomal transmembrane glycoprotein, as endogenous reporters in standard pulse/chase radiolabeling assays. As previously shown (Tazeh et al., 2009),(Koeller & Bangs, 2018), TbCatL was initially synthesized in control cells as a typical mixture of proproteins of 53 kDa (P) and 50 kDa (X), and both proteins were converted over time to the catalytically active mature enzyme form (M, 44 kDa) (Figure 3A, lanes 1-4). This pattern was also observed in TbVps24 depleted cells (Figure 3A, lanes 5-8), however significantly elevated (~13%) amounts of all TbCatL species were consistently observed in the final media fraction (Figure 3A, lanes 10 & 12). In accordance with the increase in secreted forms, we also noticed a small decrease in the extent of processing of cell-associated TbCatL (Fig. 3B). These results suggest that TbVps24 plays a relatively minor role in normal biosynthetic trafficking of soluble TbCatL to the lysosome.

**Figure 3.**
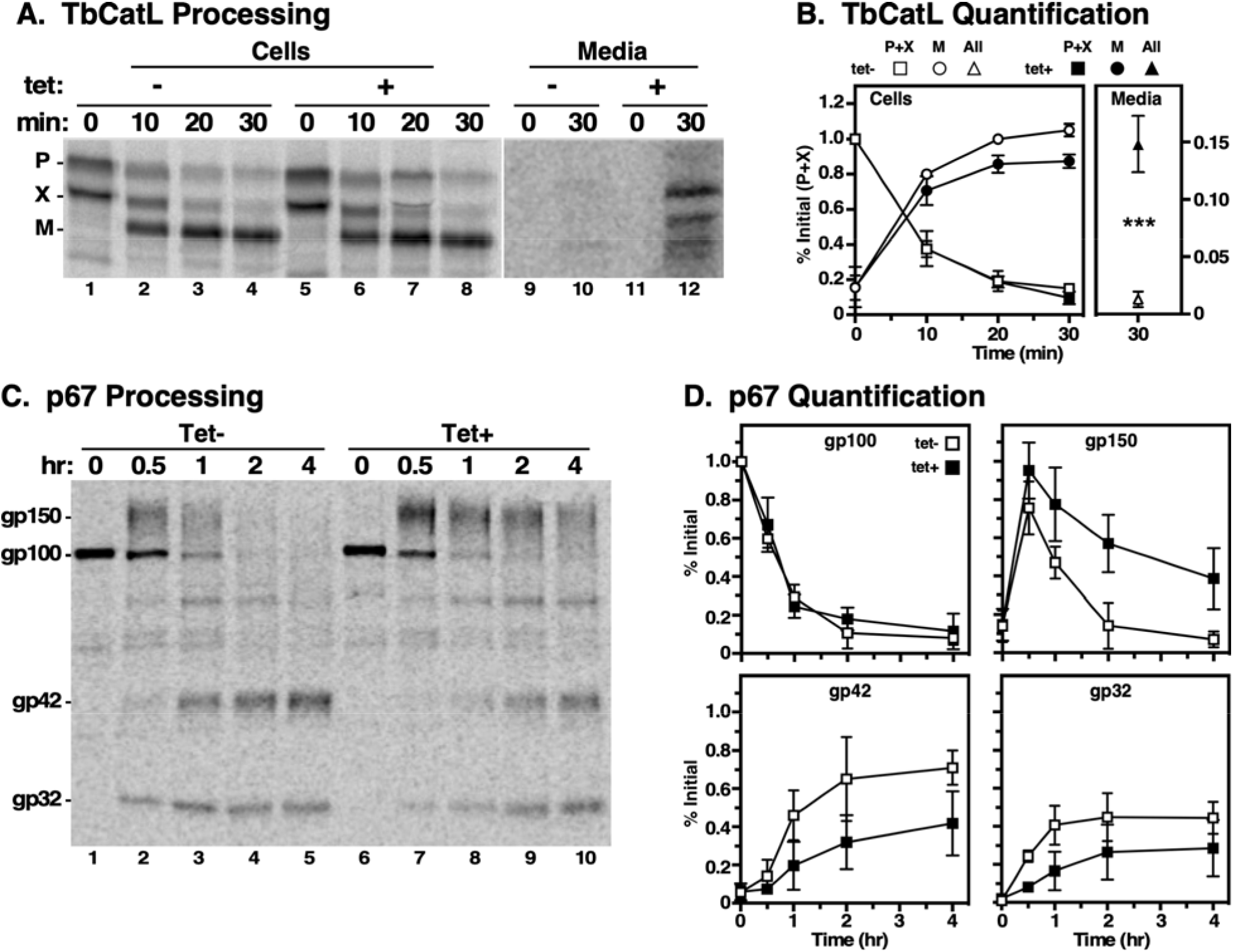
Biosynthetic trafficking to the lysosome. TbVps24 silencing was induced (12 hr) and pulse chase radiolabeling was performed followed by immunoprecipitation with anti-TbCatL (**A & B**) or anti-p67 (**C & D**) at the indicated chase time points. **A.** TbCatL samples were fractionated by SDS-PAGE and a representative phosphorimage is presented. Cell and media samples are from the same gel/image, but were digitally separated and individually contrasted to allow ready visualization of the media samples. **B**. Conversion of precursor forms (P+X) to mature TbCatL (M) was quantified and is presented as mean ± std. dev. (n=3). Recovery of media samples is offset with an expanded scale. *** p<0.001. **C.** Likewise, p67 immunoprecipitates were fractionated and a representative phosphorimage is presented. **D.** Conversion of the gp100 to gp150, and subsequently to gp42 and gp32, was quantified (mean ± std. dev., n=3).

We next analyzed trafficking of the lysosomal transmembrane glycoprotein, p67 (Alexander, Schwartz, Balber, & Bangs, 2002; Kelley, Alexander, Cowan, Balber, & Bangs, 1999). As previously reported, in control cells p67 is initially synthesized as an ER glycoform of 100 kDa (gp100), which is then converted by N-glycan modification in the Golgi to a 150 kDa glycoform (gp150) (Fig. 3C, lanes 1-3). Upon arrival in the lysosome, gp150 is cleaved into quasi-stable gp42 and gp32 glycoforms (Fig. 3C, lanes 3-5), and a similar overall pattern of processing is seen in TbVps24 silenced cells (Fig. 3C, lanes 6-10). However, quantitative analyses reveal that while silencing has no effect on disappearance of gp100, representing ER-to-Golgi transport, subsequent transport to the lysosome was dramatically delayed as shown by the decreased rate of conversion of gp150 to gp42 and gp32 (Fig. 3D). These results are very similar to the effects of TbVps4 depletion on the trafficking of TbCatL and p67 (Silverman et al., 2013). Overall then, it appears that the late ESCRT machinery is required for biosynthetic trafficking of endogenous lysosomal cargo in BSF trypanosomes, albeit more strikingly for membrane cargo than soluble.

### TbVps24 depletion accelerates VSG shedding

VSG turnover is mediated by slow shedding of VSG from the cell surface by cleavage of the glycosylphosphatidylinositol anchor with a *t*_1/2_ of ~30 hours (Bulow, Nonnengasser, & Overath, 1989; Seyfang, Mecke, & Duszenko, 1990). It has also been reported that trypanosomes release nanotubes, which form extracellular vesicles containing VSG (Szempruch et al., 2016). To investigate the role of TbVps24 in VSG shedding, we performed pulse/chase radiolabeling followed by immunoprecipitation of labeled VSG from both cell and media fractions of control and TbVps24 silenced cells (Figure 4A). In control cells <5% of total VSG was detected in the media following the 4 hr chase period, barely above the T_o_ signal (Figure 4A, lanes 3 vs 1). Under TbVps24 depletion, a significant increase (~21% at 4 hr) was observed, which comprised both full length VSG (V) and an unusual VSG fragment (X) (Figure 4A, lane 6). Inhibition of TbCatL activity blocked the production of this cleaved VSG (Figure 4C, lane 8). Thus it is likely that VSG en route to the cell surface is subject to proteolysis by TbCatL that has been misdirected into the secretory pathway by TbVps24 ablation (see Figs 3A & 3B). Previously our lab has shown that depletion of the essential housekeeping enzyme ornithine decarboxylase (similar kinetics of silencing and lethality) had no effect on VSG shedding (Umaer, Bush, & Bangs, 2018), confirming that the increase in shedding observed here is related to the knockdown of TbVps24, and not due to a non-specific effect on cell viability.

**Figure 4.**
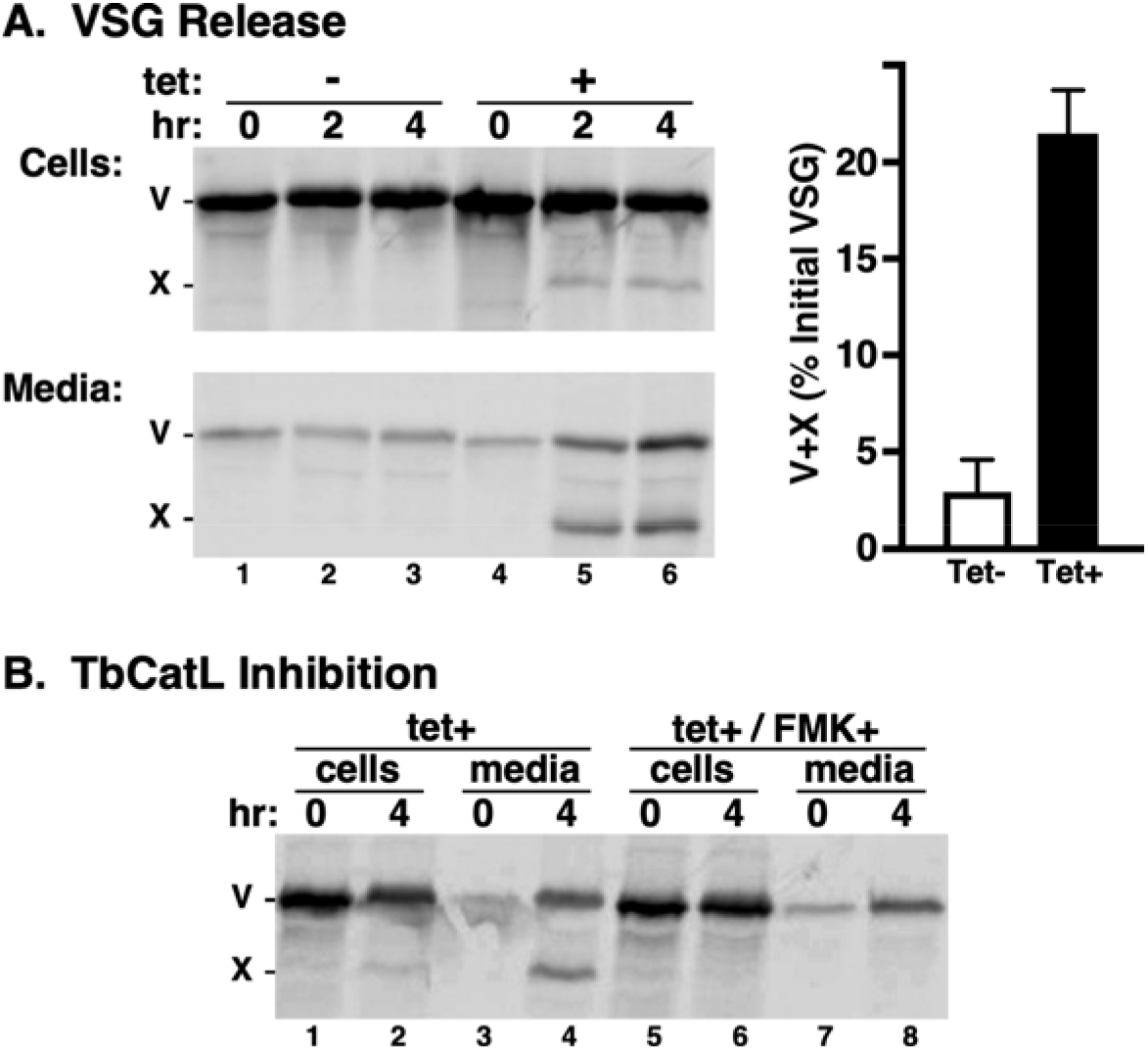
Release of VSG. **A.** TbVps24 silencing was induced (12 hr) and VSG release was monitored by pulse chase radiolabeling and immunoprecipitation from cell (top left) and media (bottom left) fractions. Both panels are from the same gel/image and were processed identically prior to digital separation for presentation. Full length (V) and proteolytic (X) species are indicated. Released VSG (V+X) was quantified (mean ± std. dev., n=3). **B.** VSG release was assayed as above in the absence or presence of 20 μM FMK024 to block endogenous TbCatL activity.

### Effect of TbVsp24 silencing on endocytosis

We next assessed the effect of TbVps24 silencing on trafficking of receptor-mediated endocytic cargo, and subsequent delivery to the lysosome, using tomato lectin (TL) and transferrin (Tf) in standard flow cytometry assays (Umaer et al., 2018). TL binds to membrane glycoproteins in the flagellar pocket, and is then taken up and delivered to the lysosome via the late endosome (Silverman et al., 2011). Tf binds to the transferrin receptor in the flagellar pocket and likewise is trafficked to the lysosome. Endocytosis of these markers was assayed under two conditions: at 4°C to allow binding without internalization and at 37°C to allow continuous uptake and lysosomal delivery. Total uptake of both TL and Tf were unimpaired by TbVps24 knockdown, nor was TL binding affected (note that this assay is not sensitive enough to measure Tf binding) (Fig. 5A). In contrast, apparent lysosomal delivery of the pH-sensitive probe TL:FITC, which is quenched as it enters progressively more acidic endosomal compartments, was significantly impaired (Fig. 5B). This could be due to two possibilities: i) the overall rate of uptake and transit of endosomal compartments is slower in silenced cells; or ii) the pH of the terminal lysosome is elevated by TbVps24 ablation. However, the fact that the curve for uptake is very similar to that of control cells, albeit offset, and that a stable end point is reached, suggests that trafficking is unaffected. To test this, the location of continuously internalized TL:biotin was determined by fluorescence microscopy (Fig. 5C). As predicted, lysosomal localization of TL was qualitatively the same in both control and silenced cells, consistent with an alteration of terminal lysosomal pH. The likely cause of this is an impact on delivery of the vacuolar ATPase (proton pump) to endolysosomal compartments.

**Fig 5.**
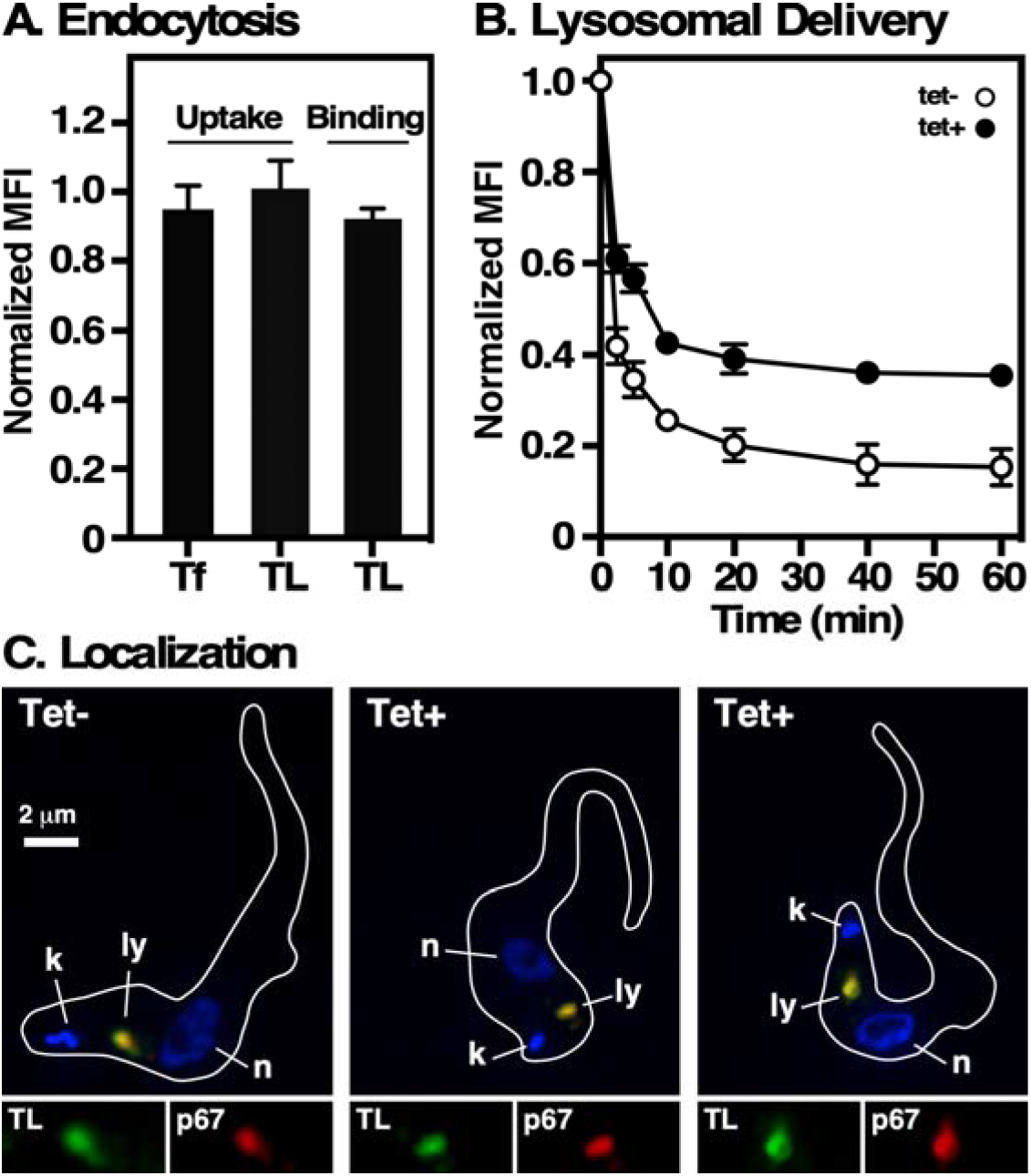
Effect of TbVps24 depletion on endocytosis. Control and TbVps24 silenced cells (12 hr) were used for ligand binding and uptake assays as described in Materials and Methods. **A.** Binding (TL:A488 only) and uptake (Tf:A488 and TL:A488) of receptor-mediated endocytic cargo were measured by flow cytometry. Mean fluorescence intensities (MFI) normalized to control un-silenced cells are presented (mean ± std. dev., n=3 biological replicates). **B.** The kinetics of endocytosis was determined by pulse-chase uptake of the pH-sensitive probe TL:FITC. Cell-associated fluorescence of live cells was measured over time by flow cytometry. Data are normalized to initial bound ligand and are presented as MFI (mean ± std. dev., n=3 biological replicates). **C.** Cells were pulsed with TL:Bio (30 min, 37°C), washed, and then reincubated (20 min, 37°C) to chase the lectin probe into terminal endocytic compartments. Imaging was performed on fixed/permeabilized cells stained with anti-p67 (red) and streptavidin:A488 (green). Representative 3-channel images are presented of control (tet-) and silenced cells (tet+). Inserts are corresponding 1.5x single channel images of the lysosomal region. Cell outlines were captured from matched DIC images. Bar indicates 2 μm.

### TbVps24 depletion affects ISG65 turnover

Knockdown of TbVsp4, the terminal ESCRT ATPase, results in accelerated lysosomal degradation of the cell surface transmembrane protein ISG65 (Silverman et al., 2013). We repeated these experiments in conjunction with TbVps24 knockdown using a standard cycloheximide chase protocol–termination of translation followed by immunoblotting to assess steady state levels of ISG65. As seen previously with TbVps4 silencing, at the end of the chase period steady state ISG65 fell to 60% of initial, and this loss was rescued by TbCatL inhibition, confirming turnover in the lysosome (Fig. 6A, lanes 1-4). Silencing of TbVps24 (Fig. 6A) resulted in both lower initial steady state levels of ISG65 (lanes 1 vs. 5) and accelerated turnover (lanes 5 & 6), the later of which is reversed by TbCatL inhibition (lanes 6 vs. 8). Next, to determine if increased degradation also impacted steady state surface levels of ISG65 we performed surface biotinylation (Fig. 6). A strong and equal signal was detected for biotinyl-VSG in both control and silenced cells (lanes 1 & 2), while no signal was detected in either for lysosomal TbCatL (lanes 3 & 4). These results confirm both plasma membrane integrity and the specificity of labeling. Likewise a strong signal was seen for surface ISG65 in control cells, but this was reduced ~80% in TbVps24 depleted cells (lanes 5&6). qRT-PCR analyses indicate that ISG65 mRNA levels, and by inference synthesis, were unaffected by TbVps24 knockdown. These results, which mimic exactly our findings with TbVps4 knockdown (Silverman et al., 2013), clearly indicate that the late ESCRT machinery in trypanosomes is involved in recycling of the internalized pool of ISG65 back to the cell surface, as opposed to delivery to the lysosome for degradation.

**Figure 6.**
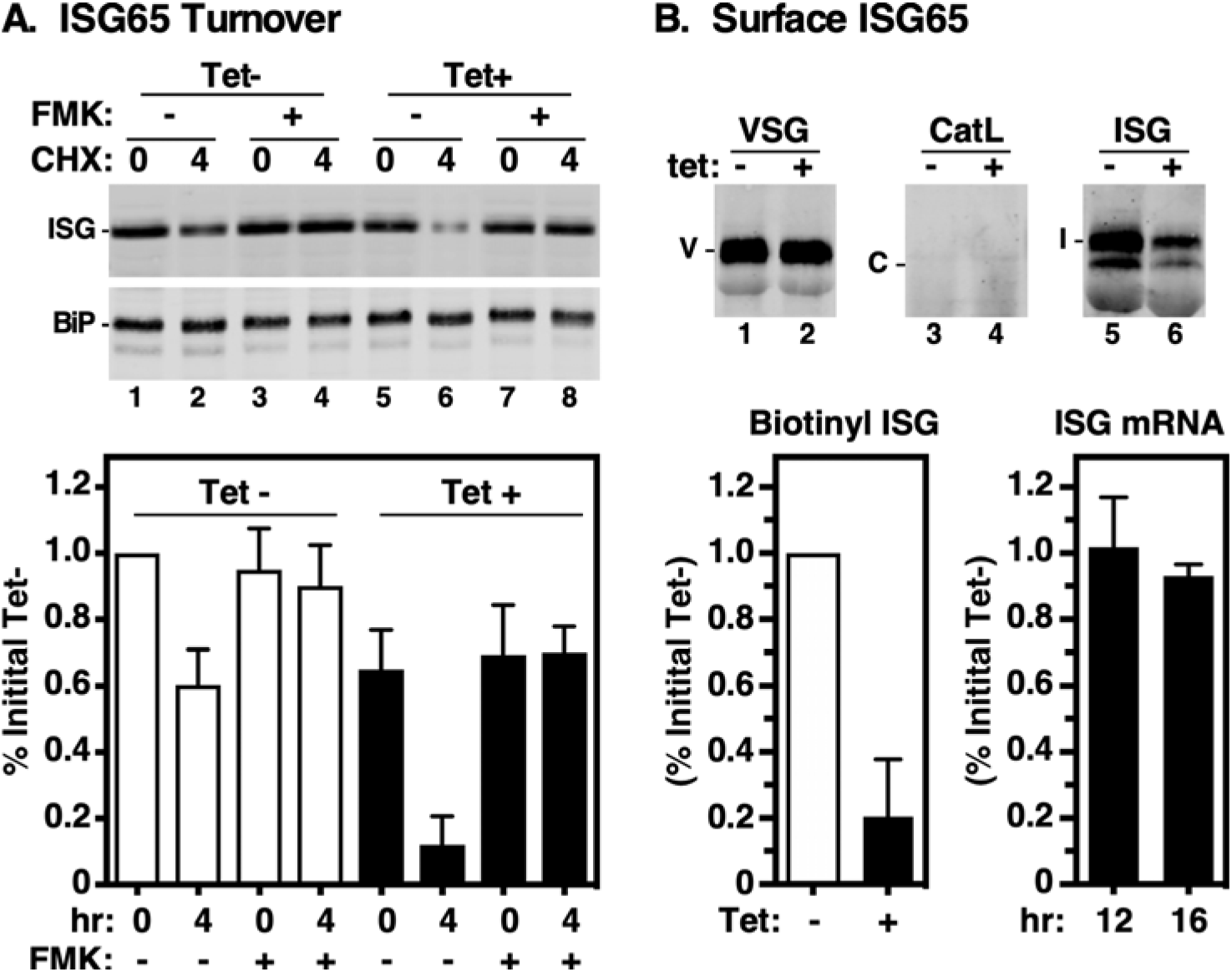
Vps24 depletion accelerates ISG65 turnover. Turnover of endogenous ISG65 was analyzed in control (Tet-) and TbVps24 silenced (Tet+) cells (12 hr). **A. Top.** Protein synthesis was inhibited by treatment with cycloheximide and steady state levels of ISG65 were assayed by immunoblotting at 0 and 4 hrs of culture. As indicated FMK024 (20 μM) was used to block degradation in the lysosome. Parallel blots were performed with anti-BiP as a loading control. Representative blots (5×10^6^ cells equivalents per lane). **Bottom.** Quantification of ISG65 levels. ISG65 signals were normalized to BiP signals and corrected values were expressed as percentage of the initial signal for uninduced cells (mean ± std. dev., n=3 biological replicates). Control cells, white bars; silenced cells black bars. **B. Top.** Intact control (Tet-) and TbVps24 silenced (Tet+) cells were surface biotinylated and immunoprecipitated with anti-VSG221 (positive control), anti-TbCatL (negative control), and anti-ISG65. After fractionation by SDS-PAGE (ISG65 & TbCatL, 10^7^ cell equivalents/lane; VSG221, 5×10^4^ cell equivalents/lane), blotting was performed with streptavidin-IR dye. Representative Li-Cor images are presented. Mobilities of VSG (V), TbCatL (C), and ISG65 (I) are indicated. Panels are from the same gel/image and were digitally separated after image processing. **Bottom Left.** ISG65 biotinylation was quantified and expressed as percentage of initial signal for uninduced cells (mean ± std. dev., n=3 biological replicates). Control cells, white bars; silenced cells black bars. **Bottom Right.** ISG65 mRNA levels were assessed by qRT-PCR at 12 (tet-) and 16 (tet+) hrs of TbVps24 silencing. Values are normalized to uninduced controls (mean ± std. dev., n=3 biological replicates)

### Recombinant TbVps24 specifically binds to phosphoinositides PI(3,5)P_2_ and PI(4,5)P_2_

Knock down of the PI(3)P-5 kinase, TbFab1, in BSF trypanosomes causes a reduction in PI(3,5)P_2_ levels and a complete blockage of ISG65 turnover in the lysosome (Gilden et al., 2017). Mammalian Vps24 and related ESCRT III components have been shown to bind PI(3,5)P_2_ and PI(P3,4)P_2_ (Catimel et al., 2008; Whitley et al., 2003). To establish a connection between the effects of TbVps24 and TbFab1 knockdown we assayed for direct binding of recombinant TbVps24 to phosphoinositides using a protein-lipid overlay dot blot approach with PIP-Strips^®^ containing 15 different phospholipids, including 7 phosphoinositides. Strong binding of the manufacturer supplied positive control (PLC-δ1 PH domain) with PI(4,5)P_2_ was seen, while no binding was observed with any lipid by recombinant TbZFP, a trypanosome RNA binding protein (Fig. 7, left and middle panels respectively). In contrast, recombinant TbVps24 bound strongly to both PI(3,5)P_2_ and PI(4,5)P_2_ (Fig. 7, right panel). No binding to any other phospholipid was seen.

**Fig 7.**
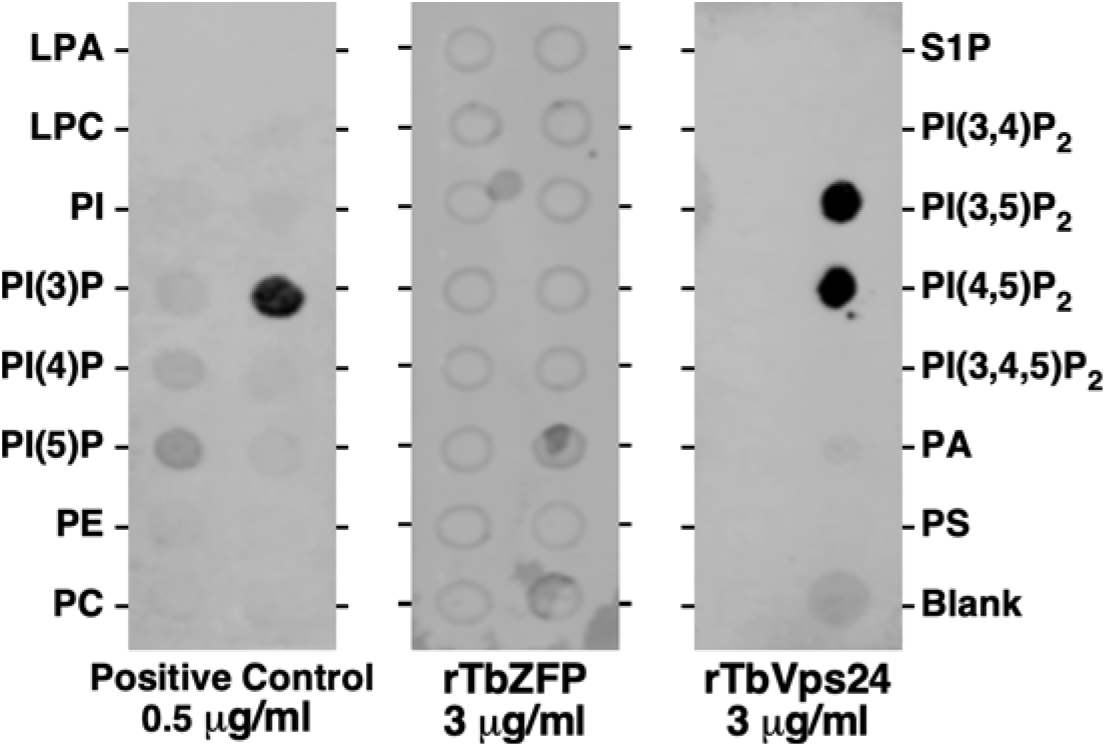
Binding to phosphoinositides. The manufacturer supplied positive control (PLC-δ1 PH domain, 0.5 μg/ml), recombinant TbZFP (negative control, 3 μg/ml) and recombinant TbVps24 (3 μg/ml) were incubated with phospholipid strips. After washing blots were probed with anti-GST (positive control) or anti-His antibody (recombinant *T. brucei* proteins). Representative Li-Cor images are presented. LPA, lysophosphatidic acid; LPC, lysophosphatidylcholine; PI, phosphatidylinositol; PE, phosphatidylethanolamine; PC, phosphatidylcholine; S1P, sphingosine-1-phosphate; PA, phosphatidic acid; PS, phosphatidylserine.

## DISCUSSION

We have performed detailed analyses of the role of TbVps24 in endomembrane trafficking in BSF trypanosomes. This ESCRT III component localizes to a Rab7^+^ late endosomal compartment that is in close proximity to the terminal p67^+^ lysosome. This same localization was seen previously for TbVps23 (ESCRT I) and TbVps4 (post-ESCRT) (Silverman et al., 2013). Silencing of TbVps24 is lethal, confirming its essentiality. General cell morphology was normal for up to 16 hrs of dsRNA induction, but cell death ensued rapidly thereafter. Consequently, all phenotypic analyses were performed at 12 hr so as to stay within the window of viability.

TbVps24 ablation impacted basal trafficking pathways in a manner similar to that seen previously with TbVps4 knockdown (Silverman et al., 2013). Biosynthetic trafficking to the lysosome was impaired. This effect was more dramatic for transmembrane p67 than for soluble TbCatL, but up to 15% of newly synthesized TbCatL was misdirected into the secretory pathway. Uptake of receptor-mediated cargo was normal, as was delivery to the lysosome. Interestingly however, the pH of the terminal lysosome was elevated, a phenotype which we attribute to impaired biosynthetic sorting of the vacuolar ATPase/H^+^ pump and/or associated regulatory proteins. This result differed from TbVps4 knockdown, in which delivery of receptor-mediated cargo to the lysosome was retarded, but the final pH was normal. TbVps24 silencing also affected normal turnover of VSG leading to an increase in shedding. It is likely that this is a secondary effect of TbVsp24 knockdown since the normal trafficking pathway of VSG to the cell surface does not involve late endosomal compartments. However, much of this shed material was proteolytically cleaved by TbCatL. The VSG fragment was also seen in the cell fraction, suggesting that VSG en route to the cell surface encounters TbCatL that has been misdirected into the secretory pathway. The effect of TbVps4 silencing on VSG turnover was not investigated (Silverman et al., 2013).

However, the most dramatic effect of TbVps24 silencing was acceleration of ISG65 turnover in the lysosome, resulting in reduced steady state levels on the cell surface. This phenotype mirrors precisely the effect of knocking down TbVps4, but contrasts with knockdown of TbVps23 (ESCRT I), which was also lethal but had no effect on ISG65 turnover (Silverman et al., 2013). These results suggest that the late ESCRT machinery, specifically, is actually involved in recycling of ISG65 from endosomal compartments back to the cell surface. It is also formally possible that enhanced degradation of ISG65 is due to the small portion of TbCatL that is misdirected into the secretory pathway, i.e. is extralysosomal. However, we feel that this is unlikely because: i) only a small portion of TbCatL is secreted, and of that an even smaller portion is actually in the active mature form; and ii) only a small portion of ISG65, both newly synthesized and recycling, would colocalize with secreted TbCatL, and then only briefly during intracellular transport. It is difficult to envision how such a mode of turnover could be more efficient than normal lysosomal degradation. The fact that newly synthesized VSG is subject to limited TbCatL-mediated degradation under similar circumstances is likely due to the relative abundance of each protein - 7×10^4^ molecules of ISG65 vs. 10^7^ copies of VSG (Ziegelbauer & Overath, 1992).

Finally, we have confirmed in another system that Vps24 is indeed a phosphoinositide-binding protein with specificity for both PI(3,5)P_2_ and PI(4,5)P_2_, in contrast to mammalian Vps23 which bound PI(3,5)P_2_ and PI(3,4)P_2_ (Whitley et al., 2003). This presents a paradox. Knockdown of TbVps24 blocks recycling of ISG65, leading to accelerated delivery to the lysosome and degradation, but knockdown of TbFab1, and hence the TbVps24 molecular interactor PI(3,5)P_2_, has the opposite effect - complete inhibition of ISG65 turnover (Gilden et al., 2017). Whether TbFab1 knockdown leads to increased recycling of ISG65 to the cell surface was not determined. Overall, this situation is somewhat reminiscent of EGFR trafficking in mammalian cells. siRNA knockdown of Vps24 and other ESCRT machinery blocks recycling of EGFR to the cell surface in HeLa cells (Baldys & Raymond, 2009), but it must be noted that other studies indicate that Vps24 knockdown decreases lysosomal degradation of EGFR, which is a more expected result for impaired ESCRT function (Raiborg, Malerod, Pedersen, & Stenmark, 2008). Nevertheless, a dominant negative form of Vps24 blocks recycling of claudin-1 from internal compartments to tight junctions (Dukes et al., 2011) indicating that a role for Vps24 in surface protein recycling is possible. In regard to PI(3,5)P_2_, pharmacological inhibition of Fab1 in HeLa cells resulted in a severe reduction in EGFR turnover, which was attributed to blockage of final MVB maturation, i.e., fusion with terminal lysosomes (de Lartigue et al., 2009). Similar effects were seen for developmental signaling receptors with *Fab1* mutations in *Drosophila* (Rusten et al., 20026), again attributed to failure to fuse with terminal lysosomes.

Synthesizing these various results allows construction of a possible roadmap for ISG65 trafficking pathways (Fig. 8), based on the assumption that PI(3,5)P_2_ is not required for the ISG65 recycling function of ESCRT, despite the proven PI(3,5)P_2_-binding properties of TbVps24. This maybe a reasonable assumption given that the late ESCRT machinery functions at many subcellular sites where PI(3,5)P_2_ is not known to localize, e.g., cellular abscission, viral budding, plasma membrane repair, and nuclear envelope maintenance, and many of these functions do not require the full armament of early ESCRT I & II components (Hurley, 2015; Schöneberg et al., 2017; Vietra et al., 2020). First, we propose that the effect of TbFab1 knockdown (inhibition of ISG65 turnover) is at the level of delivery from the LE to the terminal lysosome via the classical MVB pathway, as proposed for metazoan signaling receptors (Fig. 8, #1). The resulting depletion of PI(3,5)P_2_ would interfere with the function of TbVps24 and other ESCRT III components. Presumably the phenocopying effect of TbRab28 knockdown is also exerted at this point (Lumb, Leung, Dubois, & Field, 2011). Second, knockdown of TbRab11 blocks the recycling endosome (RE)-to-flagellar pocket pathway (Fig. 8, #2), leading to enhanced ISG65 turnover with concomitant reduction in the surface pool (Umaer et al., 2018). Under this condition ISG65 accumulates internally and presumably ‘back flows’ into degradative pathway(s). Ablation of TbRME-8 has a similar effect (Koumandou, Boehm, Horder, & Field, 2013), but not enough is known to ascribe a precise site-of-function with confidence. Third, the critical checkpoint for ISG65 fate seems to be divergence into either the pathway to the lysosome for degradation (Fig. 8, #1), as all ISG65 eventually does, or rescue and return to the cell surface via the recycling endosome, which is likely to happen a finite number of times in the lifetime of any one ISG65 molecule (Fig. 8, #3). We propose that the late ESCRT components (TbVps24 and TbVps4) are involved in both pathways. Delivery to the lysosome is dependent on ubiquitinylation of the ISG65 cytoplasmic domain (Chung et al., 2004; Chung et al., 2008; Leung et al., 2011), and since recruitment via ubiquitin recognition is a function of ESCRT I & II, we assume that the degradative pathway utilizes all of the classic ESCRT machinery. On the other hand, we suggest that a limited repertoire of late ESCRT components (and other factors?) mediate the recruitment of endosomal membrane, including ISG65, for recycling to the flagellar pocket. Finally, the critical factor in determining which fate prevails may be ISG65 ubiquitinylation. The best evidence for this is that knockdown of the early endosomal deubiquitinylase, TbVdu1, enhances ISG65 turnover by increasing the ubiquitin state of ISG65 (Zoltner, Leung, Alsford, Horn, & Field, 2015), which would be recognized by early ESCRT and recruited into the degradative pathway (Fig. 8, #4). Non-ubiquitinylated ISG65 would not be recognized and thus would be available for recycling via the late ESCRT-dependent pathway.

**Figure 8.**
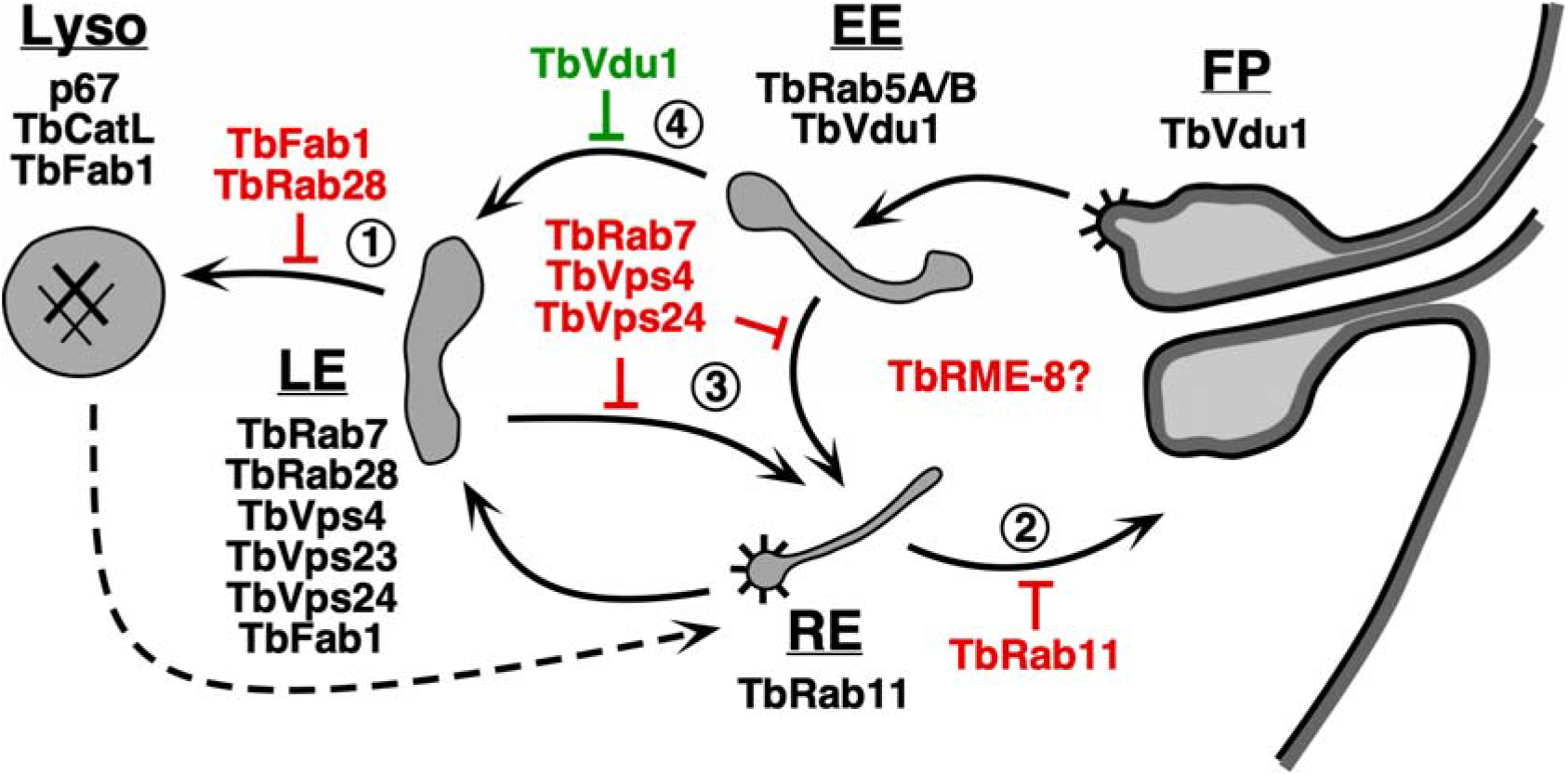
Proposed ISG65 trafficking pathways. Schematic diagram of the endocytic pathways of trypanosomes. Black arrows indicate documented routes between the flagellar pocket (FP), early endosome (EE), recycling endosome (RE), late endosome (LE), and the terminal lysosome (Lyso). Dashed arrow indicates uncertain pathway for exocytosis of degraded endocytic cargo (Hall et al., 2005). Validated markers for each compartment are: EE, TbRab5A/B (Pal, Hall, Nesbeth, Field, & Field, 2002); RE, TbRab11 (Jeffries, Morgan, & Field, 2001; Umaer et al., 2018); LE, TbRab7 (Engstler et al., 2004; Silverman et al., 2011), TbRab28 (Lumb et al., 2011), TbVps4 (Silverman et al., 2013), TbVps23 (Leung et al., 2008; Silverman et al., 2013), and TbVps24 (this work); Lyso, p67 (Alexander et al., 2002) and TbCatL (Koeller & Bangs, 2018; Peck et al., 2008). TbFab1 localizes to both LE and Lyso (Gilden et al., 2017). TbVdu1 localizes to the FP/EE region (Zoltner et al., 2015). Spiny coats indicate known sites of clathrin coated vesicle formation (Grunfelder et al., 2003; Morgan, Allen, Jeffries, Hollinshead, & Field, 2001). Points at which RNAi silencing of a given component may block an ISG65 pathway are indicated (red). The point at which TbRME-8 silencing blocks recycling of ISGs, leading to enhanced destruction, is not clear (Koumandou et al., 2013). Knockdown of TbVdu enhances ISG65 turnover (green). Numbers indicate steps referred to in the text.

This model leads to several obvious questions. First, where exactly is the divergence point? Any explanation must take into account that knockdown of the quintessential late endosomal marker, TbRab7, phenocopies loss of TbVps24 and TbVps4, i.e., ISG65 turnover is accelerated (Silverman et al., 2013). At one extreme it would be in the LE where the bulk of all documented ESCRT components, as well as TbRab7, reside in trypanosomes. At the other extreme is the EE where formation of MVBs is thought to initiate (Vietra et al., 2020), and where low levels of Rab7 are also found in conjunction with the formation of retromer complexes for recycling sorting receptors back to the Golgi (Naslavsky & Caplan, 2018). In the middle is the concept that passage from the EE to the LE is a conversion procession in which Rab5 is exchanged for Rab7, internal pH is acidified, and MVB formation matures, rather than a process of vesicular transport between static compartments (Naslavsky & Caplan, 2018). In this scenario, the divergence point could be anywhere in a continuum between the EE and LE proper. A second question is whether non-ubiquintinylated ISG65 is actively sorted into the recycling pathway, or whether this is a passive default process? Active sorting would likely require an adaptor protein as ESCRT III proteins are not known to have cargo binding properties. Thirdly, is PI(4,5)P_2_ involved in this process? This phosphoinositide is abundant in the early secretory pathway of trypanosomes (Demmel et al., 2014), as it is in all eukaryotes, and TbVsp24 binds this lipid as well as PI(3,5)P_2_. Unfortunately, testing this novel idea will be difficult as knockdown of the kinase that synthesizes PI(4,5)P_2_ blocks global endocytosis (Demmel et al., 2014), which would over-shadow any effect on recycling. A final question is if, as we propose, late ESCRT is involved in both the degradative and recycling pathways, why does knockdown of TbVps24 and TbVps4 not block ISG65 turnover as well. Why does ISG65 not stay at a fixed steady state level, or even rise due to ongoing synthesis? The answer probably lies in the fungible nature of endosomal pathways in general, and in trypanosomes in particular. When one route is blocked there is nothing to prevent a given cargo, either biosynthetic or endocytic, from flowing into another. This concept has already been invoked above to explain heightened degradation of ISG65 when TbRab11 is silenced (Fig. 8, #2). Another example is the lysosomal membrane glycoprotein p67. Biosynthetic trafficking of this protein is dependent on both clathrin and canonical di-leucine motifs in its cytoplasmic domain, but when the cytoplasmic domain is deleted it still arrives in the lysosome with similar kinetics via an alternate ‘default’ route (Alexander et al., 2002; Tazeh et al., 2009). That these routes differ is confirmed by the fact that knock down of both TbVps4 and TbRab11 dramatically reduces lysosomal trafficking of native p67, but has no effect on the truncated mutant (Silverman et al., 2013; Umaer et al., 2018). Thus it is not difficult to envision that ‘doomed’ ISG65 that normally goes to the lysosome in an ESCRT-dependent manner will still find a way when this route is blocked.

In summary, we have affected a detailed characterization of the ESCRT III subunit TbVps24 in *T. brucei*, leading to a model for the trafficking pathways of ISG65. This is of course a simplified model, but it provides a starting point for further investigation as outlined above. One additional area of interest will be to see if the same pathways hold for the distinct ISG75 family, which has been implicated as the receptor for uptake of the trypanocidal drug suramin (Alsford, Field, & Horn, 2013). Dissecting ISG75 trafficking will help elucidate the as of yet uncertain mechanism of action of this compound. Finally it should be noted that trypanosomes, which contain all the common eukaryotic pathways and compartments, albeit in simplified and streamlined format, and with a robust armament of genetic techniques, provide a powerful model system for basic cell biology investigations.

## MATERIALS AND METHODS

### Parasite maintenance and cell lines construction

Bloodstream form *T. brucei brucei* (Lister 427 strain), and its single marker (SM) derivative for tetracycline-responsive expression (Wirtz, Leal, Ochatt, & Cross, 1999), were used in all experiments. Cells were cultured in HMI-9 media (Hirumi & Hirumi, 1994) supplemented with 10% fetal bovine serum (Tet free) at 37°C and 5% CO_2_. Cells were harvested at mid-to-late log phase (0.5-1.0 x 10^6^/ml) for all experiments. A stem-loop TbVps24 dsRNA plasmid, pLew100v5x:pex11-TbVps24, was created using the entire coding sequence (660 bp) of TbVps24 (TriTrypDB:Tb927.11.10000) as described previously (Silverman et al., 2013). The construct was linearized by NotI and transfected into SM cells by electroporation (Burkard, Fragoso, & Roditi, 2007). Clonal cell lines were selected with phleomycin. Synthesis of dsRNA was induced with 1 μg/ml tetracycline. For localization experiments, TbVps24 was in situ epitope tagged at the C-terminus with 3xHA using the pXS6puro:TbVps23-3xHA plasmid (Silverman et al., 2013). The entire Vps24 orf was PCR amplified using primers with flanking 5’ HindIII and 3’ XhoI restriction sites. Digested product was used to replace the Vps23 orf with the Vps24 orf in-frame with the C-terminal 3xHA tag. Next, the Vps24 3’ UTR (nts +1 to +880 relative to the TbVps24 stop codon) was PCR amplified using primers with flanking 5’ PacI and 3’ SacI sites, and restricted product was used to replace the Vps23 3’ UTR. The final construct contains (5’–3’): TbVps24:3xHA, the aldolase intergenic region, the puromycin resistance cassette and the TbVps24 3’ UTR. All segments were confirmed by DNA sequencing. The resultant plasmid was excised with HindIII/SacI for transfection into a host cell line containing in situ Ty-tagged TbRab7 (Silverman et al., 2013; Silverman et al., 2011), and clonal cell lines were obtained under puromycin selection.

### Cloning and expression of recombinant proteins

Recombinant 6XHis-tagged TbVps24 (N-terminus tag) was cloned using the pTrcHis TOPO TA expression system (Thermo Fisher Scientific, Carlsbad, CA) as per the manufacturer’s instructions. The entire TbVps24 orf was amplified from T. brucei genomic DNA using primers TbVps24-F (5’ ATGTTTTCCGGGTGGTTCAA 3’) and TbVps24-R (5’ TCAACTAACAGTCCCACGAA 3’) and cloned into the TOPO vector. The resultant plasmid was transformed into TOP10 one-shot chemically competent E. coli cells, and expression of recombinant protein was induced with 1 mM IPTG for 3 hours at 37°C. Purification was performed using the Probond Purification System” (Thermo) as per manufacturer’s instructions. Purity was assessed by SDS-PAGE in conjunction with Coomassie blue staining and immunoblotting with anti-His antibody. Recombinant TbZFP control protein was a generous gift of Professor Laurie Read (University at Buffalo).

### Primary and secondary antibodies, and blotting reagents

The rabbit anti-VSG221, rabbit anti-BiP, rabbit anti-TbCatL, and mouse monoclonal anti□p67 antibodies have been described previously (McDowell, Ransom, & Bangs, 1998; Peck et al., 2008). Rabbit anti-ISG65 was the kind gift of Professor Mark Carrington (University of Cambridge). Rabbit anti□HA was from Sigma Aldrich. Mouse monoclonal anti-Ty was from the hybridoma Facility of the University of Alabama. Secondary antibodies include IR Dye 680 or IR Dye 800 conjugated goat anti rabbit or anti-mouse IgG (Li□Cor Biosciences, Lincoln, NE). Following SDS-PAGE, gels were transferred to Immobilon□P membranes (Millipore Corporation., Bedford, MA) using the Trans Blot Turbo apparatus (BioRad). Membranes were blocked and then incubated with primary and secondary antibodies in Odyssey Blocking Buffer (Li□Cor). Blots were visualized and fluorescent signals were scanned on an Odyssey CLx Imager (Li□Cor).

### Protein-lipid overlay assays

Identification of lipids interacting with recombinant Vps24 protein was performed using PIP-strips (Echelon Biosciences Inc., Salt Lake City, UT, USA) as per manufacturer’s instructions. The strips were blocked with 3% fatty acid-free BSA (Sigma-Aldrich) in PBS-T (PBS, 0.1% Tween-20) and then incubated (2 hr, RT) with gentle agitation in blocking buffer with 0.5-3 mg/ml of recombinant proteins. After washing with PBS-T, bound proteins were detected by immunoblotting with primary mouse anti-His (Sigma□Aldrich, Taiwan) or anti-GST primary antibodies (Sigma□Aldrich, Saint Louis, MO) and secondary goat antimouse IgG:IRDye800CW (Li-Cor).

### Quantitative real-time PCR

The levels of mRNA knockdown of TbVps24 RNAi cells were quantified using primers specific for TbVps24 (FP 5’ GATGAAGTTGAGAAGGTTGTGG 3’ and RP 5’ AACCTTGCCACTAACTCGTC 3’). RNA isolation, DNase treatment, cDNA synthesis and quantitative PCR were performed as described previously (Umaer et al., 2018). TbZFP was used as a reference gene. ISG65 mRNA levels were determined using similar methods described previously (Umaer et al., 2018).

### Endocytosis assays

The following ligand conjugates were used for endocytosis assays: transferrin:Alexa Fluor 488 (Tf:A488, Molecular Probes, Eugene, OR); tomato lectin:biotin and tomato lectin:fluorescein (TL:Bio & TL:FITC, Vector Laboratories, Burlingame, CA); and tomato lectin:Alexa Fluor 488 (TL:A488, prepared as in (Tazeh et al., 2009)). TL:Bio was detected by Streptavidin:Alexa Fluor 488 (SA:A488, Molecular Probes).

Endocytosis assays were performed as described previously (Silverman et al., 2011). Briefly, trypanosomes were washed in HBS (50 mM HepesKOH, pH 7.5, 50 mM NaCl, 5 mM KCl, 70 mM glucose) and pre-incubated (10^6^/ml) in serum-free HMI9 with BSA (0.5 mg/ml) for 10 minutes at 37°C followed by addition of tomato lectin (TL:488 or TL:FITC) or transferrin (Tf:488) for flow cytometry, or TL:Bio for epifluorescence microscopy, at a final concentrations of 5 μg/ml. For binding experiments (TL only) cells were incubated for 30 min on ice. For uptake experiments (TL and Tf) cells were incubated for 30 min at 37°C, washed and then labeled cells were analyzed by flow cytometry. For lysosomal delivery assay, TL:Bio labeled cells were incubated in fresh media for 20 minutes to chase ligand into the terminal lysosome followed by microscopy as described previously (Silverman et al., 2011). TL:Bio was detected in fixed permeabilized cells using SA:488. To determine the rate of endocytosis and lysosomal delivery cells were labeled with TL:FITC as above under binding conditions, then washed into fresh binding medium, and incubated at 37°C to internalize and chase bound ligand. Cells were then processed for flow cytometry.

### Pulse-chase radiolabeling and immunoprecipitation

Pulse-chase analyses were performed using metabolic radiolabelling with [^35^S]methionine/cysteine (Perkin Elmer, Waltham, MA, USA) and subsequent immunoprecipitation of specific radiolabeled proteins were conducted as previously described (Alexander et al., 2002; Silverman et al., 2013). Pulse times: 2 min for VSG, 10 min for TbCatL and 15 min for p67. Chase times are specified in the relevant figures. Immunoprecipitated proteins were analyzed by SDS-PAGE followed by phosphorimaging of dried gels using a Molecular Dynamics Typhoon system with native ImageQuant Software (GE Healthcare, Piscataway, NJ).

### ISG65 turnover

A cycloheximide ‘chase’ protocol was used to assay ISG65 turnover. De novo protein synthesis was inhibited by incubation with cycloheximide (100 μg/mL), and FMK024 (morpholinourea-phenylalanine-homophenylalanine-fluoromethyl ketone; 20 μM; MP Biomedicals, Aurora, OH) was used prior (15 min) to and during cycloheximide treatment to inhibit lysosomal thiol protease activity. At the indicated times, whole cell lysates were fractionated by SDS-PAGE and blotted using primary anti-ISG65 antibody and secondary goat anti-rabbit IgG:IRDye800CW (Li-Cor).

### Surface biotinylation

Biotinylation of surface proteins was performed as reported previously (Gruszynski, DeMaster, Hooper, & Bangs, 2003). Cell surface amino groups were biotinylated in PBSG by treating with membrane impermeant Sulfo-NHS-Biotin (Thermo Fisher Scientific) for 30 min on ice (1 mM final, 5×10^7^ cells/ ml). The reaction was quenched with TrisHCl, pH 7.5 (25 mM for 5 min on ice, 5×10^6^ cells/ ml). Cells were washed three times in PBSG and processed for immunoprecipitation followed by western analysis with streptavidin-IR dye 800 CW (Li-Cor).

### Epifluorescence microscopy

Immunofluorescence (IFA) microscopy was performed as previously described (Silverman et al., 2013; Silverman et al., 2011). In short, log-phase BSF parasites were fixed with 2% formaldehyde and permeablized with 0.5% NP-40 followed by blocking, incubation with primary antibodies, and stained with appropriate Alexa488-or Alexa594 conjugated secondary antibodies. Slides were washed and mounted in DAPI fluoromount G (Southern Biotech, Birmingham, AL) to reveal nuclei and kinetoplasts. Serial 0.2 micron image stacks (Z-increment) were collected with capture times from 100–500 msec (100x PlanApo, oil immersion, 1.46 numerical aperture) on a motorized Zeiss Axioimager M2 stand equipped with a rear-mounted excitation filter wheel, a triple pass (DAPI/FITC/Texas Red) emission cube, and differential interference contrast (DIC) optics. Images were captured with an Orca AG CCD camera (Hamamatsu, Bridgewater, NJ) in Volocity 6.0 acquisition software (Improvision, Lexington, MA), and individual channel stacks were deconvolved by a constrained iterative algorithm, pseudocolored, and merged using Volocity 6.1 Restoration Module. Images presented are summed stack projections of merged channels. The xyz pixel precision of this arrangement has been previously validated (Sevova & Bangs, 2009).

### Data analyses

Phosphorimages and fluorescent blot scans were quantified using ImageJ (http://imagej.nih.gov/ij/). For band intensity quantification, signals from each lane were subtracted from the signal of the equivalent unlabeled areas of that lane. All data analyses were conducted in the Prism 6 software (GraphPad Software, Inc, San Diego, CA). Replicates are indicated in the figures.

## ACKNOWLEDGMENTS

The authors are grateful to Professor Mark Carrington (Cambridge University, UK) for anti-ISG65 antibody and to Professor Laurie Read for recombinant TbZFP. This work was supported by United States Public Health Service Grants R01 AI056866 to JDB, and by support of the Jacobs School of Medicine and Biomedical Sciences.

## Notes

### Competing Interest Statement

The authors have declared no competing interest.

## REFERENCES

Alexander, D. L., Schwartz, K. J., Balber, A. E., & Bangs, J. D. (2002). Developmentally regulated trafficking of the lysosomal membrane protein p67 in Trypanosoma brucei. J. Cell. Sci., 115(Pt 16), 3253–3263.

Allen, C. L., Liao, D., Chung, W. L., & Field, M. C. (2007). Dileucine signal-dependent and AP-1-independent targeting of a lysosomal glycoprotein in *Trypanosoma brucei*. Mol Biochem Parasitol, 156(2), 175–190.

Alsford, S., Field, M. C., & Horn, D. (2013). Receptor-mediated endocytosis for drug delivery in African trypanosomes: fulfilling Paul Ehlich’s vision of chemotherapy. Trends in Parasitology, 29, 207–212.

Baldys, A., & Raymond, J. R. (2009). Critical role of ESCRT machinery in EGFR recycling. Biochemistry, 48(40), 9321–9323.

Balla, T. (2013). Phosphoinositides: tiny lipids with giant impact on cell regulation. Physiol Rev, 93(3), 1019–1137.

Bangs, J. D. (2011). Replication of the ERES:Golgi junction in bloodstream-form African trypanosomes. Mol Microbiol, 82(6), 1433–1443.

Behnia, R., & Munro, S. (2005). Organelle identity and the signposts for membrane traffic. Nature, 438(7068), 597–604.

Bulow, R., Nonnengasser, C., & Overath, P. (1989). Release of the variant glycoprotein during differentiation of bloodstream to procyclic forms of *Trypanosoma brucei*. Mol. Biochem. Parasitol., 32, 85–92.

Burkard, G., Fragoso, C. M., & Roditi, I. (2007). Highly efficient stable transformation of bloodstream forms of *Trypanosoma brucei*. Mol. Biochem. Parasitol., 153(2), 220–223.

Catimel, B., Schieber, C., Condron, M., Patsiouras, H., Connolly, L., Catimel, J., … Holmes, A. (2008). The PI(3,5)P2 and PI(4,5)P2 interactomes. Journal of Proteome Research, 7, 5295–5313.

Chung, W. L., Carrington, M., & Field, M. C. (2004). Cytoplasmic targeting signals in transmembrane invariant surface glycoproteins of trypanosomes. Journal of Biological Chemistry, 279(52), 54887–54895.

Chung, W. L., Leung, K. F., Carrington, M., & Field, M. C. (2008). Ubiquitylation is required for degradation of transmembrane surface proteins in trypanosomes. Traffic, 9(10), 1681–1697.

de Lartigue, J., Polson, H., Feldman, M., Shokat, K., Tooze, S. A., Urbe, S., & Clague, M. J. (2009). PIKfyve regulation of endosome linked pathways. Traffic, 10, 883–893.

Demmel, L., Schmidt, K., Lucast, L., Havicek, K., Zankel, A., Koestler, T., … Warren, G. (2014). The endocytic activity of the flagellor pocket in *Trypanosoma brucei* is regulated by an adjacent phosphatidylinositol phosphate kinase. Journal of Cell Science, 127, 2351–2364.

Di Paolo, G., & De Camilli, P. (2006). Phosphoinositides in cell regulation and membrane dynamics. Nature, 443(7112), 651–657.

Dukes, J. D., Fish, L., Richardson, J. D., Blaikley, E., Burns, S., Caunt, C. J., … Whitley, P. (2011). Functional ESCRT machinery is required for constitutive recycling of claudin-1 and maintenence of polarity in vertebrate epithelial cells. Molecular Biology of the Cell, 22, 3192–3205.

Engstler, M., Bangs, J. D., & Field, M. C. (2006). Intracellular transport systems in trypanosomes: function, evolution and virulence. In M. J. Barry JD, McCulloch R, Acosta-Serrano A (Ed.), Trypanosomes – After the Genome (pp. 281–317). Wymondham, UK: Horizon Scientific Press.

Engstler, M., Thilo, L., Weise, F., Grünfelder, C. G., Schwarz, H., Boshart, M., & Overath, P. (2004). Kinetics of endocytosis and recycling of the GPI-anchored variant surface glycoprotein in *Trypanosoma brucei*. J. Cell. Sci., 117, 1105–1115.

Gilden, J. K., Umaer, K., Kruzel, E. K., Hecht, O., Correa, R. O., Mansfield, J. M., & Bangs, J. D. (2017). The role of the PI(3,5)P2 kinase TbFab1 in endo/lysosomal trafficking in *Trypanosoma brucei*. Mol. Biochem. Parasitol., 214, 52–61.

Grunfelder, C. G., Engstler, M., Weise, F., Schwarz, H., Stierhof, Y. D., Morgan, G. W., … Overath, P. (2003). Endocytosis of a glycosylphosphatidylinositol-anchored protein via clathrin-coated vesicles, sorting by default in endosomes, and exocytosis via RAB11-positive carriers. Mol. Biol. Cell., 14(5), 2029–2040.

Gruszynski, A. E., DeMaster, A., Hooper, N. M., & Bangs, J. D. (2003). Surface coat remodeling during differentiation of *Trypanosoma brucei*. Journal of Biological Chemistry, 278(27), 24665–24672.

Hall, B. S., Smith, E., Langer, W., Jacobs, L. A., Goulding, D., & Field, M. C. (2005). Developmental variation in Rab11-dependent trafficking in *Trypanosoma brucei*. Eukaryot. Cell., 4(5), 971–980. Retrieved from

Hirumi, H., & Hirumi, K. (1994). Axenic culture of African trypanosome bloodstream forms. Parasitol. Today, 10(2), 80–84.

Hurley, J. H. (2015). ESCRTs are everywhere. European Journal of Biochemistry, 34, 2398–2407.

Jeffries, T. R., Morgan, G. W., & Field, M. C. (2001). A developmentally regulated rab11 homologue in *Trypanosoma brucei* is involved in recycling processes. J. Cell. Sci., 114(Pt 14), 2617–2626.

Kabiri, M., & Sterverding, D. (2000). Studies on the recycling of the transferrin receptor in *Trypanosoma brucei* using an inducible gene expression system. European Journal of Biochemistry, 267, 3309–3314.

Kelley, R. J., Alexander, D. L., Cowan, C., Balber, A. E., & Bangs, J. D. (1999). Molecular cloning of p67, a lysosomal membrane glycoprotein from *Trypanosoma brucei*. Mol. Biochem. Parasitol., 98(1), 17–28.

Koeller, C. M., & Bangs, J. D. (2018). Processing and targeting of cathepsin L (TbCatL) to the lysosome in Trypanosoma brucei. Celluar Microbiology, e12980.

Koumandou, V. L., Boehm, C., Horder, K. A., & Field, M. C. (2013). Evidence for recycling of invariant surface transmembrane domain proteins in African trypanosomes. Eukaryot. Cell., 12(2), 330–342.

Leung, K. F., Dacks, J. B., & Field, M. C. (2008). Evolution of the multivesicular body ESCRT machinery; retention across the eukaryotic lineage. Traffic, 9(10), 1698–1716.

Leung, K. F., Riley, F. S., Carrington, M., & Field, M. C. (2011). Ubiquitylation and developmental regulation of invariant surface protein expression in trypanosomes. Eukaryot. Cell., 10(7), 916–931.

Lumb, J. H., Leung, K. F., Dubois, K. N., & Field, M. C. (2011). Rab28 function in trypansoomes: interactions with retromer and ESCRT pathways. Journal of Cell Science, 124, 3771–3783.

McDowell, M. A., Ransom, D. M., & Bangs, J. D. (1998). Glycosylphosphatidylinositol-dependent secretory transport in *Trypanosoma brucei*. Biochem. J., 335 (Pt 3), 681–689.

Morgan, G. W., Allen, C. L., Jeffries, T. R., Hollinshead, M., & Field, M. C. (2001). Developmental and morphological regulation of clathrin-mediated endocytosis in *Trypanosoma brucei*. J. Cell. Sci., 114(Pt 14), 2605–2615.

Naslavsky, N., & Caplan, S. (2018). The enigmatic endosome-sorting the ins and outs of endocytic trafficking. J. Cell. Sci., 131, jcs.216499.

Overath, P., & Engstler, M. (2004). Endocytosis, membrane recycling and sorting of GPI-anchored proteins: *Trypanosoma brucei* as a model system. Mol. Micro., 53, 735–744.

Pal, A., Hall, B. S., Nesbeth, D. N., Field, H. I., & Field, M. C. (2002). Differential endocytic functions of *Trypanosoma brucei* Rab5 isoforms reveal a glycosylphosphatidylinositol-specific endosomal pathway. Journal of Biological Chemistry, 277(11), 9529–9539. \

Peck, R. F., Shiflett, A. M., Schwartz, K. J., McCann, A., Hajduk, S. L., & Bangs, J. D. (2008). The LAMP-like protein p67 plays an essential role in the lysosome of African trypanosomes. Mol. Microbiol., 68(4), 933–946.

Raiborg, C., Malerod, L., Pedersen, N. M., & Stenmark, H. (2008). Differential functions of Hrs and ESCRT proteins in endocytic membrane trafficking. Experimental Cell Research, 314, 801–813.

Rusten, T. E., Rodahl, L. M. W., Pattni, K., Englund, C., Samakovlis, C., Dove, S., … Stenmark, H. (20026). Fab1 phosphatidylinositole 3-phosphate kinase controls trafficking but not silencing of endocytosed receptors. Molecular Biology of the Cell, 17, 3989–4001.

Schöneberg, J., Lee, I.-H., Iwasa, J. H., & Hurley, J. H. (2017). Reverse-topology membrane scission by the ESCRT proteins. Nat. Rev. Mol. Cell. Biol., 18.

Sevova, E. S., & Bangs, J. D. (2009). Streamlined architecture and glycosylphosphatidylinositol-dependent trafficking in the early secretory pathway of African trypanosomes. Mol. Biol. Cell., 20(22), 4739–4750.

Seyfang, A., Mecke, D., & Duszenko, M. (1990). Degradation, recycling, and shedding of *Trypanosoma brucei* variant surface glycoprotein. J Protozool, 37(6), 546–552.

Silverman, J. S., & Bangs, J. D. (2012). Form and function in the trypanosomal secretory pathway. Curr. Opin. Microbiol., 15(4), 463–468.

Silverman, J. S., Muratore, K. A., & Bangs, J. D. (2013). Characterization of the late endosomal ESCRT machinery in *Trypanosoma brucei*. Traffic, 14(10), 1078–1090.

Silverman, J. S., Schwartz, K. J., Hajduk, S. L., & Bangs, J. D. (2011). Late endosomal Rab7 regulates lysosomal trafficking of endocytic but not biosynthetic cargo in *Trypanosoma brucei*. Mol. Microbiol., 82(3), 664–678.

Szempruch, A. J., Sykes, S. E., Kieft, R., Dennison, L., Becker, A. C., Gartrell, A., … Harrington, J. M. (2016). Extracellular Vesicles from *Trypanosoma brucei* Mediate Virulence Factor Transfer and Cause Host Anemia. Cell, 164(1-2), 246–257.

Tazeh, N. N., Silverman, J. S., Schwartz, K. J., Sevova, E. S., Sutterwala, S. S., & Bangs, J. D. (2009). Role of AP-1 in developmentally regulated lysosomal trafficking in *Trypanosoma brucei*. Eukaryot. Cell., 8(9), 1352–1361.

Umaer, K., Bush, P. J., & Bangs, J. D. (2018). Rab11 mediates selective recycling and endocytic trafficking in *Trypanosoma brucei*. Traffic, 19(6), 406–420.

Vietra, M., Radulovic, M., & Stenmark, H. (2020). The many functions of ESCRTs. Nature Reviews Molecular Cell Biology, 25–41.

Whitley, P., Reaves, B. J., Hashimoto, M., Riley, A. M., Potter, B. V., & Holman, G. D. (2003). Identification of mammalian Vps24p as an effector of phosphatidylinositol 3,5-bisphosphate-dependent endosome compartmentalization. J Biol Chem, 278(40), 38786–38795.

Wirtz, E., Leal, S., Ochatt, C., & Cross, G. A. (1999). A tightly regulated inducible expression system for conditional gene knock-outs and dominant-negative genetics in *Trypanosoma brucei*. Mol. Biochem. Parasitol., 99(1), 89–101.

Ziegelbauer, K., & Overath, P. (1992). Identification of invariant surface glycoproteins in the bloodstream stage of *Trypanosoma brucei*. Journal of Biological Chemistry, 267, 10791–10796.

Zoltner, M., Leung, K. F., Alsford, S., Horn, D., & Field, M. C. (2015). Modulation of the surface proteome through multiple ubiquitylation pathways in African trypanosomes. PLoS Pathog., 11, e1005236.

